# Identifying genomic adaptation to local climate using a mechanistic evolutionary model

**DOI:** 10.1101/2024.11.21.624744

**Authors:** Nikunj Goel, Christen M. Bossu, Justin J. Van Ee, Erika Zavaleta, Kristen C. Ruegg, Mevin B. Hooten

## Abstract

1. Identifying genomic adaptation is key to understanding species’ evolutionary responses to environmental changes. However, current methods to identify adaptive variation have two major limitations. First, when estimating genetic variation, most methods do not account for observational uncertainty in genetic data because of finite sampling and missing genotypes. Second, many current methods use phenomenological models to partition genetic variation into adaptive and non-adaptive components.
2. We address these limitations by developing a hierarchical Bayesian model that explicitly accounts for observational uncertainty and underlying evolutionary processes. The first layer of the hierarchy is the data model that captures observational uncertainty by probabilistically linking RAD-sequence data to genetic variation. The second layer is a process model that represents how evolutionary forces, such as local adaptation, mutation, migration, and drift, maintain genetic variation. The third layer is the parameter model, which incorporates our knowledge about biological processes. For example, because most loci in the genome are expected to be neutral, the environmental sensitivity coefficients are assigned a regularized prior centered at zero. Together, the three models provide a rigorous probabilistic framework to identify local adaptation in wild organisms.
3. Analysis of simulated RAD-seq data shows that our statistical model can reliably infer adaptive genetic variation. To show the real-world applicability of our method, we re-analyzed RAD-seq data (∼105k SNPs) from Willow Flycatchers (*Empidonax traillii*) in the USA. We found 30 genes close to loci that showed a statistically significant association with temperature seasonality. Gene ontology suggests that several of these genes play a crucial role in egg mineralization, feather development, and the ability to withstand extreme temperatures.
4. Moreover, the data and process models can be modified to accommodate a wide range of genetic datasets (*e.g.*, pool and low coverage genome sequencing) and demographic histories (*e.g.,* range shifts) to study climatic adaptation in a wide range of natural systems.

## Introduction

Many species face a looming threat of extinction as global temperature rises (Thomas *et al*. 2004; Urban 2015). This risk is particularly concerning for species with limited dispersal capacity that are slow to track their climatic niches (Carroll *et al*. 2015). Alternatively, some species may adapt to the local climate, potentially alleviating the risk of extinction. Natural selection favors individuals with heritable phenotypes best suited to survive and reproduce in local climates, creating a geographical mosaic in frequencies of adaptive alleles congruent with climatic gradients (Hedrick, Ginevan & Ewing 1976; Hedrick 1986; Hedrick 2006). Therefore, to understand the evolutionary responses of a species to a changing climate and incorporate this knowledge to inform evolutionary management policies (Smith *et al*. 2014), we need robust methodologies to identify genes (and their function) that confer local adaptation in natural populations.

With recent advances in high-throughput sequencing and global environmental sensing technologies (Chuvieco 2020; Satam *et al*. 2023), new computational approaches have enabled researchers to identify adaptive genetic variation in wild populations. These approaches, collectively known as environmental association analysis (Rellstab *et al*. 2015; Hoban *et al*. 2016), have gained widespread popularity, which can be partly attributed to declining sequencing costs, improvements in genomic tools, detailed global climatic maps, and availability of computational resources. These advances have allowed researchers to study climatic adaptation in non-model organisms at a fine genomic resolution and large spatial extent for a reasonable cost (Frichot *et al*. 2013; De Villemereuil & Gaggiotti 2015; Fitzpatrick & Keller 2015; Wagner, Chávez-Pesqueira & Forester 2017).

Broadly, environmental association analysis involves three steps. First, sequencing data from spatially referenced individuals are used to estimate spatial variation in allele frequencies. Next, a statistical model is used to partition this genetic variation into adaptive and non-adaptive components. The adaptive variation corresponds to the response of an allele to changes in prevailing climatic conditions. The non-adaptive variation describes the allelic variation (or neutral variation) that stems from evolutionary forces other than local adaptation, including gene flow among populations, mutation, and drift. Finally, a statistical criterion is used to select loci that exhibit strong adaptive genetic variation relative to non-adaptive variation.

Despite the success of environmental association studies, current methods face several conceptual and statistical challenges that limit the reliability of the inferences. Many commonly used environmental association methods do not account for uncertainty in estimates of allele frequencies (see Coop *et al*. (2010) and Foll and Gaggiotti (2008) for exceptions), which can introduce biases in downstream analyses. Due to practical constraints, biogeographers collect and sequence only a finite number of individuals in a population. The genotype of these individuals, inferred from RAD (restriction site-associated DNA) sequencing (Baird *et al*. 2008), for instance, contains imprecise information regarding the underlying allele frequencies that result from sampling a small subset of individuals within a population. Additionally, because of low coverage and alignment errors, the true genotypes of some individuals may be missing (Huang & Knowles 2016), which further reduces sample size and increases uncertainty. Failure to account for these sources of uncertainty results in overly precise estimates of genetic variation, which can lead to an increased number of false positives (i.e., incorrectly inferring non-adaptive loci as important). Alternatively, researchers discarding data due to noisy estimates of genetic variation may lose valuable (although imprecise) information, thereby increasing the rates of false negatives (i.e., inability to identify an adaptive locus).

Another major difficulty is that many environmental association methods characterize genetic variation using models, such as generalized linear models (Rellstab *et al*. 2015). Although the underlying relationships between data and parameters (e.g., environment and allele frequency) in phenomenological models are based on evolutionary thinking, these relationships are often weakly motivated by evolutionary theory (but not always; see De Villemereuil and Gaggiotti (2015) and Günther and Coop (2013) for exceptions). For example, some classical environmental association models assume an S-shape response curve to relate environmental variables and allele frequency (Joost *et al*. 2007; Stucki *et al*. 2017). This response curve has several attractive characteristics: it is bounded between zero and one, and has an incline in the middle that emulates the response of an allele to local climatic adaptation. However, the S-shaped response curve is not unique in these characteristics. A biogeographer can construct alternate equally plausible response curves (*e.g.*, cdf functions) and, consequently, there is no easy way to choose one over the other *a priori*.

In contrast, mechanistic models of evolution are based on the first principles of birth-death processes, and, therefore, a biogeographer can evaluate model assumptions based on the constraints imposed by the population demography and natural history of the species (Rice 2004). Mechanistic models of evolution in statistical inferences offers a unique advantage—they constrain the flow of information from data to model parameters using a theoretical understanding. This may allow biogeographers to create bespoke statistical models tailored to match the demographic history of the species (Wikle 2003; Hooten & Hefley 2019).

Over the past two decades, theoretical and computational advances in Bayesian statistics have fueled a surge in the development of probabilistic hierarchical models (Beaumont & Rannala 2004; Stadler *et al*. 2024), offering researchers new avenues to address the aforementioned challenges. These hierarchical models can account for uncertainty in estimates of genetic variation (stemming from finite sampling and sequencing) (Gompert & Buerkle 2011) and then propagate this uncertainty downstream to mechanistic process models of evolution, enabling a wide range of inferences ranging from constructing phylogenies (Suchard *et al*. 2018) to identifying migration corridors (Hanks & Hooten 2013; Marcus *et al*. 2021) to estimating growth rates of emerging viral diseases (Stadler 2010). In fact, these statistical insights are also beginning to permeate the latest generation of genotype-environment association methods (Coop *et al*. 2010; De Villemereuil & Gaggiotti 2015).

Hierarchical models help us do joint inferences that allows us to learn parameters using data, process, and parameter models (Berliner 1996). In the context of environmental association analysis, the data model describes how the unobserved allele frequencies could have led to genetic data. The process model describes the biological processes determining spatial variation in allele frequencies. The parameter model describes the knowledge the biogeographer has about the parameters before the data are collected, based on past research.

This structure of the hierarchical model offers several advantages in identifying genomic adaptation to climate. First, the data model enables a biogeographer to quantify observational uncertainty in estimates of genetic variation, particularly in cases where sample sizes are small and genotypes are missing. Second, these imprecise estimates of genetic variation can then be linked to a process model of evolution. This evolution model is based on the demographic history of the species and provides a mechanistic explanation of how to partition genetic variation into adaptive (response curve) and non-adaptive (neutral) components. Lastly, the parameter model can be used to incorporate knowledge from past research. For example, the theory of molecular evolution suggests that the overwhelming majority of variation in genomes is non-adaptive and stems from the interplay between mutation, drift, and migration, while only a small fraction of the variation stems from local adaptation (Kimura 1983). To incorporate this prior knowledge, a biogeographer can assign a low prior probability that the response curve has a finite slope. This prior effectively shrinks (regularizes) the adaptive genetic variation to zero for most loci, providing a systematic way to evaluate the relative contribution of climate adaptation and neutral processes in determining genetic variation.

Because of these features of the data, process, and parameter models, hierarchical models provide a rigorous probabilistic framework to identify genomic adaptation in wild organisms informed by theoretical principles of evolutionary biology using noisy genetic data. In what follows, we present a new demographic Bayesian model and test its robustness by analyzing synthetic data with known parameter values. We use the model to analyze RAD-seq data from Willow Flycatchers (*Empidonax traillii*) and identify candidate genes that may be responsible for local climatic adaptation. Although we apply our model to Willow Flycatchers, the statistical insights are general and broadly applicable to other species.

### Willow Flycatchers

The Willow Flycatcher is a migratory songbird that breeds mainly in the United States and Southern Canada. The species is categorized into four distinct subspecies—the Pacific Northwestern form (*E. t. brewsteri*), Western Central form (*E. t. adastus*), Eastern form (*E. t. traillii*), and Southwestern form (*E. t. extimus*). Among them, the Southwestern form has experienced a precipitous decline in abundance, likely due to the loss of riparian habitat along streams and waterways (Sedgwick 2000), which provide respite during extreme temperatures. In 1995, the Southwestern form was listed as federally endangered subspecies (Unitt 1987) due to its genetic, ecological, and song distinctiveness (Theimer *et al*. 2016; Ruegg *et al*. 2018; Mahoney *et al*. 2020).

Previous landscape genomic work identified highly significant correlations between allele frequencies in genes linked to thermal tolerance and the intensity of summer heat waves in the southwest (Ruegg *et al*. 2018). Therefore, re-analyzing the Willow Flycatcher genome-wide genetic dataset, initially examined using traditional phenomenological models, offers an ideal opportunity to apply a new environmental association analysis that mechanistically accounts for genetic variation. More broadly, the results have important implications for implementing management practices designed to improve the genetic health of the endangered subspecies.

### Data

A total of 175 Willow flycatchers were sampled at 23 one-degree squares across the continental United States (Fig. 1). The sampling effort varied from 2-21 individuals per location. DNA was extracted from blood and tissue samples using the QiagenTM DNeasy Blood and Tissue extraction kit and quantified using the Qubit® dsDNA HS Assay kit (Thermo Fisher Scientific). Sequencing was conducted across three lanes of 100 bp paired-end reads on an Illumina HiSeq 2500 at the UC Davis Genome Center. We filtered SNPs using the tradeoff between discarding SNPs with low coverage and discarding individuals with missing genotypes using the R package genoscapeRtools (Anderson 2019), resulting in approximately 105k SNPs, of which 3 percent of the loci had a missing genotype (Ruegg *et al*. 2018). At each sampling location, climate data were obtained from WorldClim, which averaged the climate between 1960 and 1990 (Hijmans *et al*. 2005). Due to the high correlation between some of the top-ranked climatic variables identified in Ruegg *et al*. (2018), we limited our analysis to the four least correlated climate variables (standardized) to reduce collinearity: BIO 4 (temperature seasonality), BIO 5 (maximum temperature of the warmest month), BIO 11 (mean temperature of the coldest quarter), and BIO 17 (Precipitation in the driest quarter). For more details about the sampling design, sequencing, and bioinformatic analysis, refer to Ruegg *et al*. (2018).

**Figure 1:**
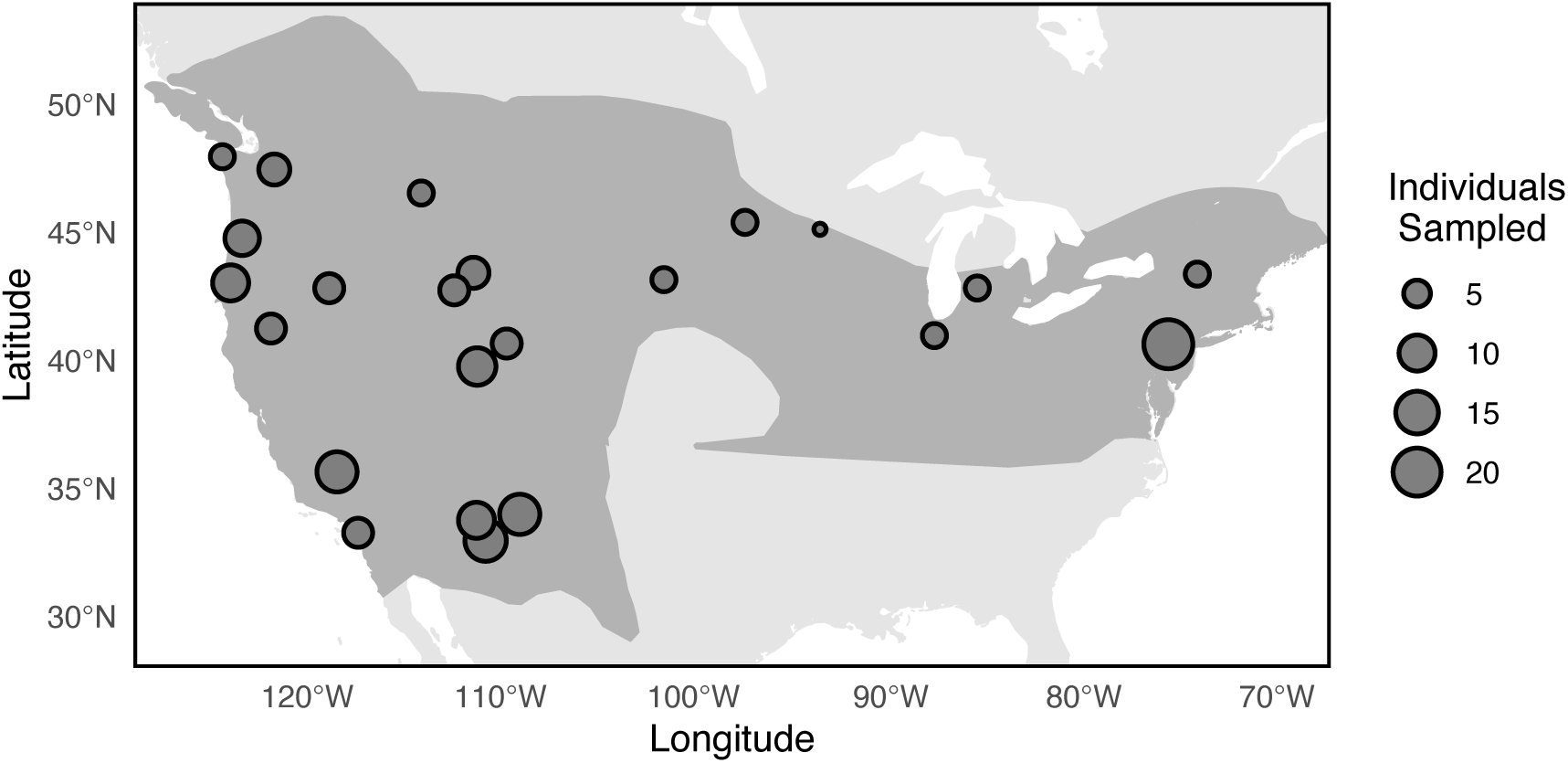
Breeding range of Willow Flycatchers (dark grey region) and 23 sampling locations. The size of the points is proportional to the number of sampled individuals in our study, which varied between 2 and 21 individuals.

### Model Formulation

To identify putative loci under selection due to local adaptation, we specified a Bayesian model with three hierarchical levels corresponding to the data, process, and parameter models. These three levels are organized such that the output of the parameter model is the input for the process model, whose output is the input for the observer model (Pagel & Schurr 2012). To facilitate a broad understanding of how the statistical model works, Figure 2 illustrates the three levels of the hierarchical model, showing how these levels are connected. In Figure 3, we provide a Directed Acyclic Graph (DAG) to illustrate the relationship between data and parameters. In the following section, we give details on the assumptions underlying the model’s construction (also see the Supplementary Information for a concise summary).

**Figure 2:**
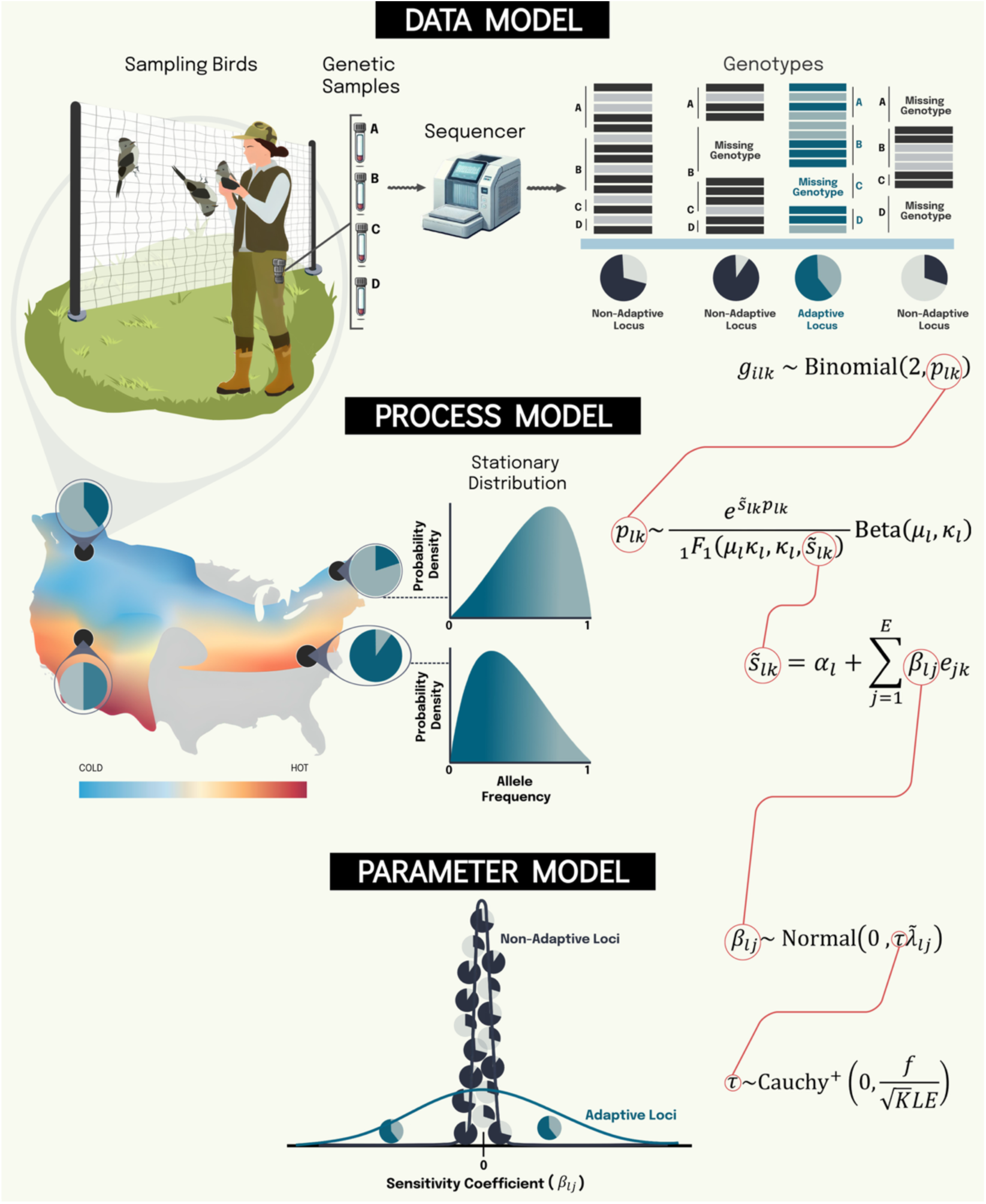
A conceptual diagram describing the hierarchical structure of the Bayesian model. The data model (*top*) links the individuals’ genotype (*g*_*ilk*_) to allele frequency (*p*_*lk*_) at a locus *l* in patch *k* (Eq. 3). The process model (*middle*) describes the stationary distribution of allele frequency resulting from evolutionary forces, such as mutation, migration, drift, and local adaptation. (Eq. 8 and 11). Finally, the parameter model (*bottom*) allows us to regularize sensitivity coefficients (β_*l*j_) using prior knowledge from molecular evolutionary theory (Eqs. [14], [15], and [16]).

**Figure 3:**
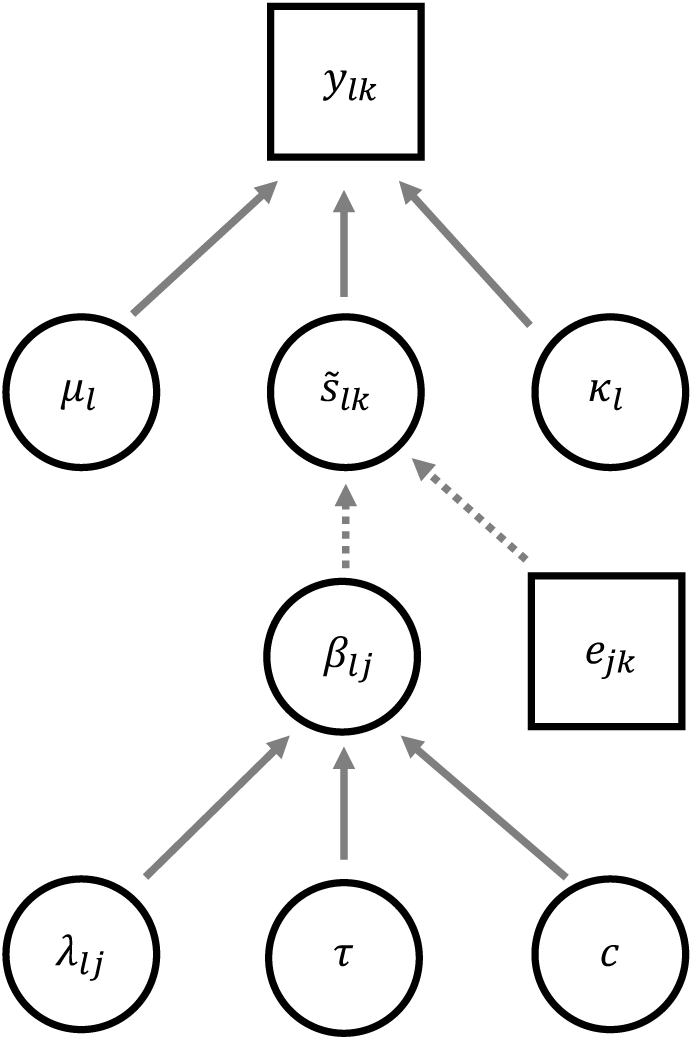
Directed Acyclic Graph (DAG) of the hierarchical Bayesian model. The squares represent data (genotype counts and environmental variables), the circles represent parameters, the dashed arrows represent deterministic relationships, and the solid arrows represent stochastic relationships.

### Data model

We assume that Willow Flycatchers were sampled at *K* sites across the United States, and each bird was genotyped at *L* independent genetic (RAD) markers (see sampling and sequencing above). We characterized the genotype of the *i*th sampled bird at site *k* using *g*_*ilk*_, which corresponds to the number of copies of the reference allele at locus *l*. Assuming individuals have biallelic loci, *g*_*ilk*_ is a discrete variable that takes values zero, one, or two. Using these individual genotype data, we define a population-level genotype variable

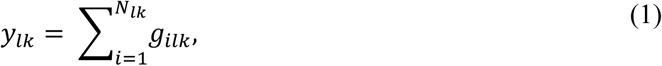

that corresponds to the number of reference alleles for *N*_*lk*_ birds that were genotyped at locus *l*. In the data, *N*_*lk*_ is always less than or equal to the number of birds sampled at a site because, for some individuals, the genotype at a locus may be missing due to sequencing errors or low coverage. Assuming the birds were randomly sampled, we model the variation in reference allele frequency (*p*_*lk*_) using a binomial distribution,

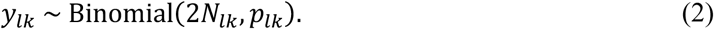

Alternatively, we can also use

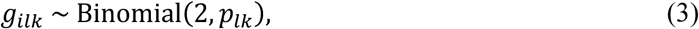

as a data model to describe the statistical relationship between allele frequency and an individual’s genotype. However, because of computational efficiency, we use equation (2). Note that the observer model accounts for the uncertainty in genetic variation that stems from finite sampling and missing genotypes.

### Process model

We consider an island-metapopulation model proposed by Wright (1931) to model evolutionary dynamics. We rely on this model because it provides a parsimonious explanation of how evolutionary processes maintain genetic variation and allows for reasonable statistical inferences even when the process model is misspecified due to complexity in evolutionary dynamics (see the Model Testing section below). We assume that the demes in the metapopulation correspond to the sites where the birds were sampled. Each deme has a population size of *N*_*e*_, and the migration rate between any pair of demes is equal. We assume that the variation in allele frequencies across demes is maintained by directional and non-directional evolutionary forces. Directional forces—such as local adaptation to climate, mutation, and migration—result in changes in the mean value of allele frequency. Non-directional forces—such as genetic drift—do not change the mean value of allele frequency but create sampling variance due to finite population size. Because of this variance, the frequency of an allele cannot be determined exactly. Instead, allele frequency is characterized probabilistically using a probability distribution, ψ(*p*_*lk*_, *t*) (Rice 2004; Blanquart, Gandon & Nuismer 2012). Using the Fokker-Planck equation, one can show that the probability distribution of the frequency of the reference allele changes as follows:

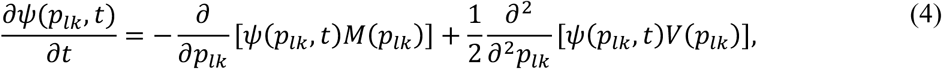

where

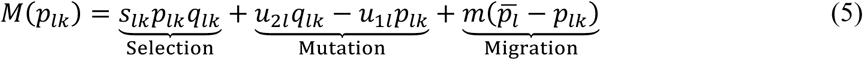

is the rate of directional change in the allele frequency and

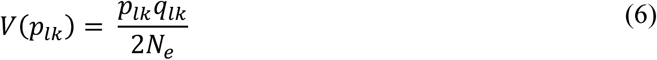

is the variance in the allele frequency due to non-directional (*i.e.*, drift) effects. In equations (4)-(6), *q*_*lk*_(= 1 − *p*_*lk*_) is the frequency of the alternate allele, *s*_*lk*_ is the environmentally regulated selection coefficient, *u*_2*l*_ and *u*_1*l*_ are the locus-specific forward and backward mutation rates, *m* is the migration rate, and *p*L_*l*_ is the average allele frequency of the immigrants. We consider multiplicative selection dynamics (*i.e.*, no dominance), such that the relative fitness of individuals with genotype (*g*_*ilk*_) zero, one, and two is 1 + 2*s*_*lk*_, 1 + *s*_*lk*_, and 1, respectively. Our model can also be modified to accommodate alternate forms of selection dynamics, such as frequency-dependent selection, underdominance, and heterozygote advantage.

At the time of sampling, the allele frequency distribution is assumed to be in quasi-equilibrium with climate conditions. To obtain this equilibrium distribution, also known as the stationary distribution (ψ_*s*_), we set ∂ψ(*p*_*lk*_, *t*)/∂*t* = 0. The stationary distribution of the allele frequency can be expressed in terms of evolutionary parameters as follows (Blanquart, Gandon & Nuismer 2012):

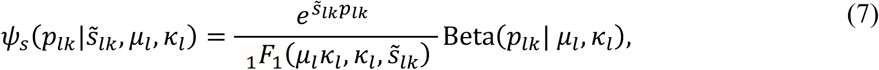

where ŝ_*lk*_ = 4*N*_*e*_*s*_*lk*_, _1_F_1_ is the hypergeometric confluent function, and μ_*l*_ = (*u*_2*l*_ + *mp*L_*l*_)/(*u*_1*l*_ + *u*_2*l*_ + *m*) and K_*l*_ = 4*N*_*e*_(*u*_1*l*_ + *u*_2*l*_ + *m*) are the mean and precision parameters of the beta distribution. To incorporate climate adaptation, we assume that the selection coefficient, ŝ_*lk*_, is regulated by local environmental conditions. For simplicity, we assume a linear relationship between the selection coefficient and *E* standardized environmental variables:

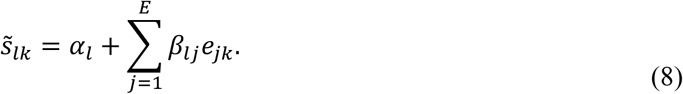

However, based on the natural history of the species, a biogeographer may consider alternative functional forms to describe the relationship between the selection coefficient and environmental variables. In equation (8), *e*_j*k*_ is the *j*th environmental variable (standardized) in deme *k*, β_*l*j_is the selection coefficient’s sensitivity to variation in the *j*th environmental variable, and α_*l*_ can be interpreted either as the selection coefficient at mean environmental conditions or a fixed value of selection coefficient arising from adaptive processes other than local climatic adaptation.

The stationary distribution in equation (7) has several notable features. The response curve of an adaptive allele (the relationship between climate and mean allele frequency),

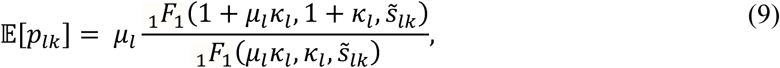

is bounded between zero and one, and its shape is determined by evolutionary parameters with biological interpretation (Blanquart, Gandon & Nuismer 2012). When selection is absent, the beta distribution captures the non-adaptive genetic variation,

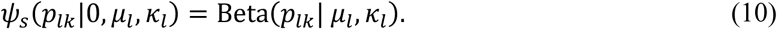

Therefore, to account for the joint contribution of adaptive and non-adaptive evolutionary forces in determining genetic variation, we use the stationary distribution to statistically model evolutionary dynamics,

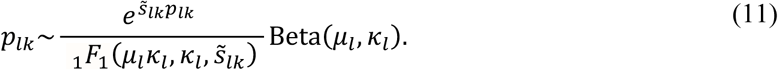

For fast and memory-efficient implementation of the statistical model, we combine the data (Eq. 2) and process (Eq. 11) models by marginalizing over *p*_*lk*_,

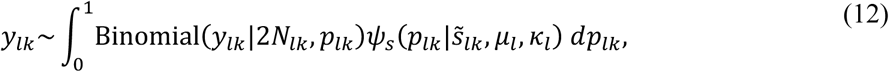

which results in an integrated likelihood that relates genotype counts to evolutionary dynamics as follows:

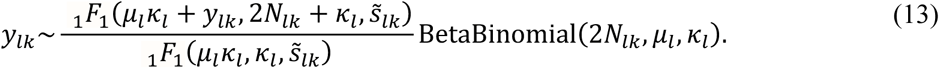

Note that the integrated likelihood in equation (13) accounts for the uncertainty that stems from noisy genetic data and the stochastic nature of evolutionary dynamics.

### Parameter model

Finally, we assign priors to parameters in equation (13) based on our prior scientific knowledge. We assign μ_*l*_ ∼ Uniform(0,1) and K_*l*_∼Normal^+^(0, 5) priors in equation (13). Due to degeneracy of the likelihood (Eq. 11, see Fig. S1), the mean of the beta distribution (μ_*l*_) and selection coefficient at the mean environment (α_*l*_) are non-identifiable (see Supplementary Information for more details). We alleviate this by fixing α_*l*_ = 0 and re-interpret μ_*l*_ as the mean of the baseline allele frequency distribution (or neutral distribution) resulting from evolutionary forces other than adaptation to variation in local climate around the mean. Next, because we expect that most of the genetic variation arises due to neutral processes (Kimura 1983), we use a regularized horseshoe shrinkage parameter model (Piironen & Vehtari 2017) for sensitivity coefficients, β_*l*j_. The horseshoe prior shrinks most of the β_*l*j_ to zero, allowing a fraction of the coefficients to take non-zero values. The horseshoe model achieves this using global (τ) and local (λe*l*j) shrinkage parameters and is expressed as

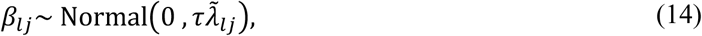

where

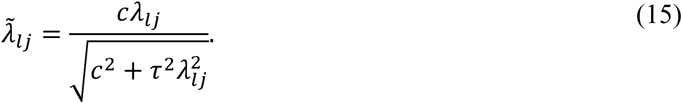

To control global shrinkage, we specify a half-Cauchy prior,

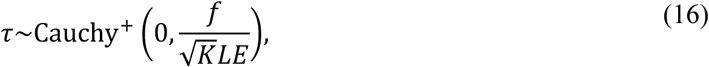

where *f* is a free variable, which we set to 20. Based on the recommendations of Piironen and Vehtari (2017), we tested a range of values of *f* and found that the statistical results are robust to orders of magnitude of variation in *f* (see Model Testing section below). Note that the global sparsity (controlled by τ) increases with the number of SNPs (*L*) and environmental variables (*E*). On the other hand, the local shrinkage parameter, λe*l*j, allows some coefficients to escape shrinkage because of a heavy-tail prior, λ_*l*j_∼Cauchy^+^(0,1). However, these large-valued coefficients are weakly shrunk with an implied regularization of Normal(0, *c*) with *c*^2^ ∼ InvGamma(2,4). We adopt an inverse-gamma prior because the prior means for both *c*^2^ (which controls adaptive dynamics via sensitivity coefficients) and K_*l*_(which governs neutral dynamics) are similar in magnitude, but the heavy right tail of the inverse-gamma distribution allows adaptive loci to attain large sensitivity coefficients. These features of the local and global shrinkage parameters effectively induce a spike and slab prior (Mitchell & Beauchamp 1988), which prevent numerical pathologies when large effect loci are weakly identifiable.

### Model Testing

To assess the robustness of our statistical method, we fit the model to synthetic data that are similar in characteristics to the Willow Flycatcher genomic data, and the generative process only partially resembles the process model considered in our statistical model. This simulation allows us to test the sensitivity of our method to model misspecifications and test its applicability to biologically realistic datasets.

We generated the synthetic genotype data in three stages (see Supplementary code for simulation details). In the first stage, we simulated selection dynamics. We considered diploid individuals with a genome size of one thousand independent loci. Each locus in the genome was assigned a random non-zero α_*l*_. We randomly selected fifteen loci that contribute to local adaptation. Each of the fifteen loci was randomly paired with one of the four environmental variables, and the corresponding selection coefficient (*s*_*lk*_) was calculated using equation (8). We used real environmental values (standardized) to calculate *s*_*lk*_to preserve the correlation structure between environmental variables.

In the second stage, we simulated metapopulation dynamics with demes equal to the number of sampling sites in the real data. We sampled random variables from the stationary distribution of allele frequencies by simulating the following stochastic differential equation (Korolev *et al*. 2010):

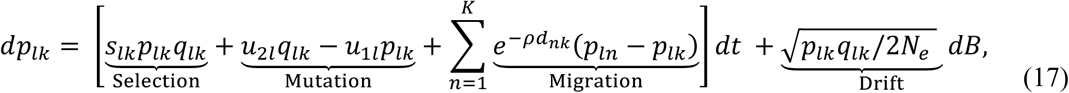

where *s*_*lk*_ is the selection coefficient obtained from stage one, *B* is the standard Brownian motion, *d*_*nk*_ is the distance between demes *n* and *k*, and ρ^−1^ is the dispersal length scale. The above equation is an Itô representation of the Fokker-Planck equation presented earlier (Eq. 4), with a small modification: migration rate depends on the distance between demes (Fig. S2).

In the third stage, we simulated genotypes. In each deme, we simulated genotypes for the same number of individuals as in the real data. For each individual, we generated a genotype at a locus by sampling from the distribution Binomial(2, *p*_*lk*_) (Eq. 3), where *p*_*lk*_ is the reference allele frequency obtained from the second stage. We randomly selected three percent of the loci and treated them as missing. To test if the statistical model can identify loci that contribute to local adaptation in synthetic data, we fit the model using Stan programming language (Carpenter *et al*. 2017) to obtain a posterior sample of sensitivity coefficients, β_*l*j_, using Hamiltonian Monte Carlo (Neal 2011). For each pair of loci and environmental variables, we computed the probability that the posterior distribution of the sensitivity coefficient included zero. If this probability is less than a threshold value of 0.05 (*p*_*t*ℎ_), we inferred that the locus may have contributed to local adaptation for the corresponding environmental variable.

To conduct robustness checks, we generated ten synthetic datasets for nine parameter regimes (a total of 90 datasets; see Supplementary Information for parameter values used to generate synthetic datasets). These parameter combinations represent three mutation and migration regimes—high, medium, and low—relative to local adaptation. For each regime, we calculated the false-negative and false-positive rates. We found that our model performs best in the low mutation-migration regime, where the genetic variation is more sensitive to climatic variation (Tables S1 and S2). However, this regime also yields higher rates of false positives. In contrast, in the high mutation-migration regime, neutral evolutionary forces play a larger role, and consequently, the statistical model results in high false negative rates.

For illustration, the Manhattan plot in Fig. 4A shows the result from a simulation corresponding to the low mutation and medium migration regime. The y-axis corresponds to the negative log probability that the posterior distribution of sensitivity coefficients included zero (Wang *et al*. 2022). The points above *y* = 2.99 (the black line) have *p*_*t*ℎ_ < 0.05. These points are characterized by a ring and solid center. The color of the center represents the true environmental variable associated with the loci, and the color of the ring represents the statistically inferred adaptive environmental variable. Note that, in this simulation run, the model incorrectly identified the corresponding environmental variable for two of the nine loci (points above the black line with a green ring and purple center). This feature can be explained by a negative correlation (−0.72) between environmental variables, BIO 4 (green) and BIO 11 (purple). As a result, the statistical model used one of the environmental variables as a substitute for another (Rellstab *et al*. 2015). We also re-analyzed the data in Fig. 4A by varying *f* over four orders of magnitude (2^0^ to 2^14^). We find that our results are nearly identical to results in Fig. 4A (see Fig. S3 and S4), suggesting that our analysis is robust to a wide variation in choice of priors for τ.

**Figure 4:**
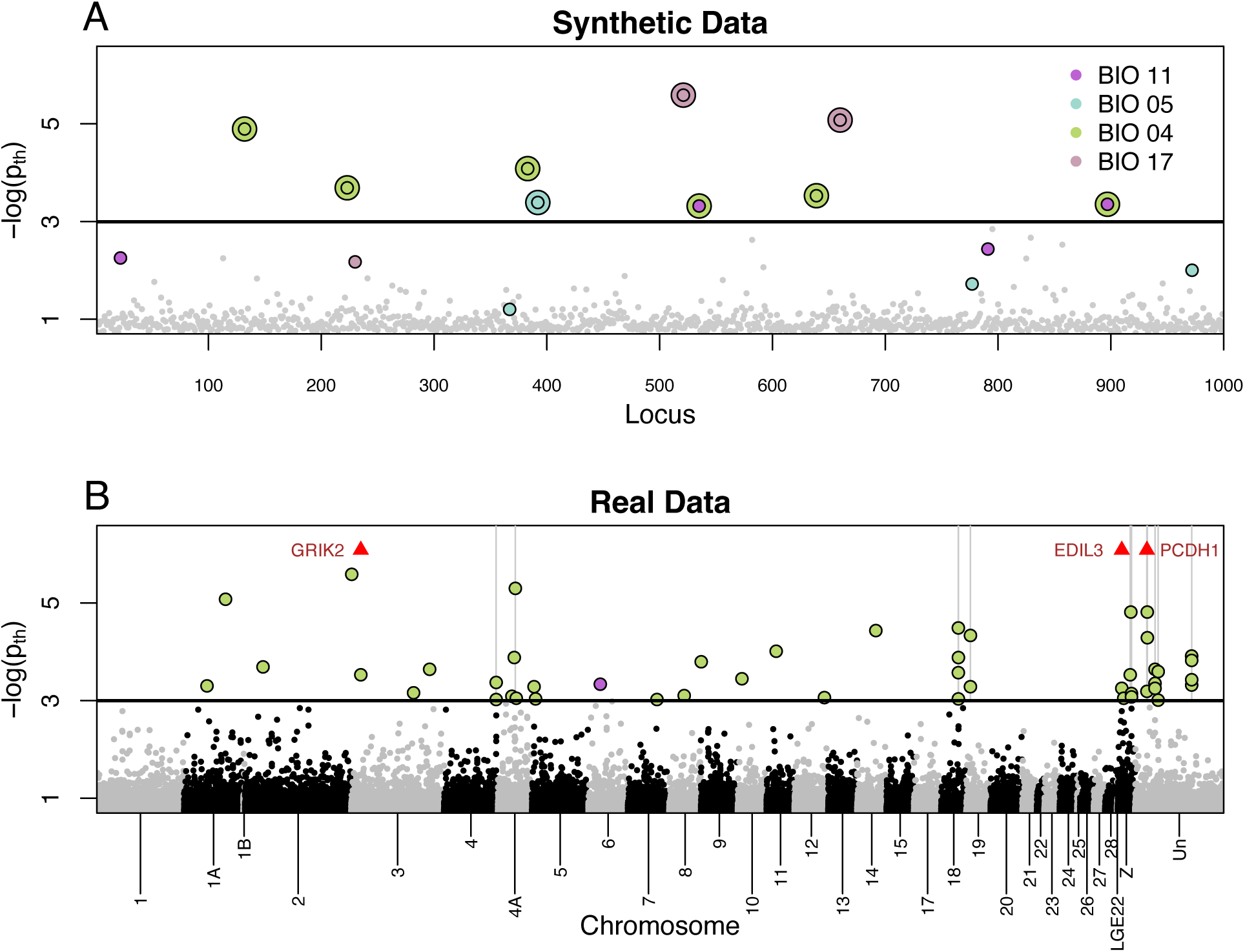
Manhattan plot showing negative log probability (− log(*p*_*t*ℎ_)) that the posterior distribution of sensitivity coefficients (β_*li*_) includes zero for synthetic (A) and real data (B). In both plots, points above the black horizontal line have *p*_*t*ℎ_ less than 0.05. (A) For synthetic data, we denote points above the black line using an outer ring and a center. The color of the center (ring) corresponds to the true (statistically inferred) environmental variable responsible for local adaptation. (B) For real data, we denote points above the black line with a solid center, with its color corresponding to the statistically inferred environmental variable responsible for local adaptation (also see Fig. S5). The vertical gray lines represent physically linked loci with statistically significant sensitivity coefficients (see Supplementary Information for parameter values used to simulate synthetic data in Fig. 4A)

Next, we re-analyzed the synthetic datasets using latent factor mixed model (LFMM; Frichot *et al*. 2013) which was used to analyze Willow Flycatcher data in Ruegg *et al*. (2018). We conducted simulations using two versions of the synthetic dataset. In the first version, we used individual genotypes, and, in the second version, we used raw allele frequencies (*i.e.*, *p*_*lk*_ = 0.5 *y*_*lk*_/*N*_*lk*_) as input. Our results suggest that LFMM yielded much higher false-negative and false-positive rates (Tables S3-S6).

### Data Analysis

Analyzing genomic data from Willow Flycatchers revealed 47 significant loci, all except one were associated with BIO 4 (Fig. 4B and Fig. S5). A closer inspection of these loci suggests that many of them cluster together on the genome (indicated by vertical gray lines in Fig. 4B). These clustering patterns are unlikely to happen by random chance. Alternatively, the observed clustering patterns can be explained by gene hitchhiking (Smith & Haigh 1974). The allele frequencies at a locus under selection and its neighboring neutral loci rise or fall in unison due to physical linkage on the chromosome. Consequently, when a gene is under selection, neutral polymorphic sites close to the gene experience environmentally regulated pseudo-selection (Barton 1998).

Using the annotated genome of Willow Flycatchers, 36 of the candidate loci were found within or nearby (within 25kb) 30 named genes (Table S7). Twenty-three of these genes are characterized, and 20 have functional roles in chicken (*Gallus gallus*) that span 8 gene ontology categories. The majority of genes cluster in 4 categories: 5 genes are involved in catalytic activity, 5 genes have binding functionality, 3 have transcription regulatory activity, and 3 have transporter activity. A closer investigation into several genes shows that EDIL3 plays a role in the egg mineralization process in Aves (Le Roy *et al*. 2021), PCDH1 is involved in feather development (Lin, Wang & Redies 2013), and GRIK2 acts as a thermoreceptor conferring sensitivity to cold temperatures in mice (Cai *et al*. 2024).

The above analysis required 32 hours to complete on a machine with a 24-core 3.6 GHz processor. By subsampling the genomic data, we also estimated the model run time for *E* = 4 and *L* = 2^8^, 2^9^, …, and 2^17^ (Fig. S6).

## Discussion

Adaptation to climate is ubiquitous and will continue to play a major role in maintaining biodiversity in the Anthropocene (Thompson 2013). However, many commonly used methods for identifying genomic adaptation are limited because they often lack the ability to handle noisy genetic data and do not formally account for the demographic history of the species. Our mechanistic Bayesian model addresses these limitations. The important aspects of our approach comprise the following features.

First, we proposed a data model that probabilistically links RAD-seq data to genetic variation (Eq. 2). This probabilistic link allowed us to quantify uncertainty in genetic variation that arises due to finite sampling and missing genotypes. Consequently, the data model properly accounts for uncertainty in the estimation of genetic variation and faithfully propagates the available information in raw genetic data to the process model for downstream analysis (Hobbs & Hooten 2025).

Second, we used a metapopulation process model to partition estimated genetic variation into adaptive and non-adaptive components (Blanquart, Gandon & Nuismer 2012). The metapopulation model can be implemented using a beta distribution to characterize neutral variation (Eq. 10) and hypergeometric confluent functions to model adaptive variation (Eq. 9). Our process model is motivated by theoretical principles from evolutionary biology. Thus, we can evaluate the model based on the underlying assumptions.

Finally, previous work shows that most of the genetic variation in wild populations is maintained by non-adaptive evolutionary forces (Kimura 1983). We incorporated this prior knowledge into our statistical model using a regularized horseshoe parameter model, which shrinks most of the selection coefficients to zero. The level of shrinkage is controlled by a global shrinkage parameter that introduces sparsity in sensitivity coefficients (Piironen & Vehtari 2017) (Eq. 14). Thus, the parameter model allowed us to partition genetic variation into adaptive (driven by environmental variation) and non-adaptive (driven by neutral processes) components.

To test our statistical model, we conducted simulations to assess the model’s performance on ninety synthetic RAD-seq datasets generated by simulating bird genomes of 1000 loci (Eq. 17), which emulate the characteristics of genetic data from Willow Flycatchers. Our statistical method performed best in the low mutation-migration regime, and the performance decreased in the high mutation-migration regime (Table S1 and S2). These results are consistent with our theoretical expectations. When the mutation and migration rates are low, genetic variation is more sensitive to variation in climate. However, in the high mutation-migration regime, neutral processes play a bigger role in maintaining genetic variation. As a result, it is more challenging to identify weakly adaptive loci.

Figure 4A shows results from a simulation from the low mutation and medium migration regime. The model correctly identified nine out of fifteen adaptive loci. However, we found that out of the nine correctly identified loci, two loci were incorrectly paired with their corresponding environmental variable due to a −0.72 correlation between BIO 4 and BIO 11. This highlights that biogeographers may need to exercise caution when interpreting statistical results or use uncorrelated predictors. The statistically inferred environmental variable might be correlated with the true environmental variable responsible for local adaptation (Rellstab *et al*. 2015). Nevertheless, the synthetic simulations suggest that our statistical model can be used with genetic datasets from wild populations.

Indeed, we identified 30 genes located within the 25 kb region flanking 47 significant loci in RAD-seq data from Willow Flycatchers, most of which were associated with temperature seasonality (Fig. 4B and S5). Some of these genes include EDIL3, PCDH1, and GRIK2, which play a role in the egg mineralization process (Le Roy *et al*. 2021), feather development (Lin, Wang & Redies 2013), and act as a thermoreceptor conferring sensitivity to cold temperatures (Cai *et al*. 2024), respectively. This suggests that temperature fluctuations may be a key driver of local adaptation in the species, providing evidence that standing genetic variation in Willow Flycatchers could alleviate or buffer extinction risk due to increasing temperature variability predicted by climate projections (Olonscheck *et al*. 2021).

Although these genes differ from those identified in Ruegg *et al*. (2018), these differences are not surprising because our hierarchical model is fundamentally different from traditional environmental association analysis in terms of quantifying uncertainty while estimating genetic variation and using an evolutionary process model to explain sources of genetic variation. Our synthetic data simulations confirm that these differences may play an important role in inferences; re-analyzing synthetic data using LFMM resulted in substantially higher rates of false positives and false negatives (Table S3-S6).

In addition to the conceptual advantages offered by our mechanistic statistical model, synthetic simulations highlight several practical scenarios where our hierarchical model may be better suited to analyze genomic data. Some of these scenarios include low sample size (between 1-4 individuals), wide dispersion in the number of individuals sampled at various locations, and a large fraction of missing genotypes. We can address these data scenarios by quantifying genetic variation probabilistically and propagating the corresponding uncertainty in downstream analyses. However, there are some computational tradeoffs: our statistical analysis required 32 hours to analyze 105k SNPs, and, as a result, our approach may require substantial computational resources to analyze genomic datasets with millions of SNPs.

Despite these computational requirements, the hierarchical structure of the model provides new opportunities to improve statistical inferences. For example, the hierarchical model is flexible and can integrate a wide range of process and data models, allowing biogeographers to better understand the evolutionary responses of a species to a changing climate while dramatically reducing the cost of analysis. The cost of sequencing varies along three axes—the number of sequenced sites in the genome, the depth of sequencing effort, and the number of sampled individuals. Typically, in RAD sequencing, a small proportion of sites in the genome are sequenced deeply to identify true genotypes, which are subsequently used to estimate allele frequencies (Baird *et al*. 2008). Although RAD sequencing is a cost-effective protocol to obtain a reduced representation of the genome, it fails to characterize a large proportion of genetic variation in the genome that could be potentially adaptive (Lowry *et al*. 2017). New sequencing protocols, such as low-depth whole genome sequencing (Alex Buerkle & Gompert 2013) and pool sequencing (Gautier *et al*. 2013), are emerging as attractive alternatives because they provide a wider genomic coverage without increasing cost or sacrificing statistical power. The key idea behind these approaches is to redistribute the same resources to sequence a larger sample of individuals and a greater proportion of the genome (Lou *et al*. 2021). This reduction in sequencing effort (per individual per locus) increases uncertainty in the estimates of individual genotypes. But, instead of discarding SNPs due to low coverage, one can model the true genotype as a parameter and propagate the corresponding uncertainty to inform population-level allele frequencies. In effect, these approaches sacrifice certainty in individual genotypes to gain genomic coverage while keeping the cost and genetic information constant. To leverage information from these sequencing protocols, the data model in the Bayesian hierarchy can be modified to account for genotype uncertainty in estimating allele frequencies.

Another potential avenue to improve inference is to construct alternate process models incorporating a wider range of demographic histories to correct for model misspecification. For example, most species, including Willow Flycatchers, are geographically structured, and, as such, the migration rates depend on the distance between demes. Closer demes exchange more migrants than demes that are far apart. This may create genetic patterns, such as isolation by distance (Fig. S2) and genetic swamping, that cannot be adequately captured by an unstructured metapopulation model (Eq. 11). Because of these misspecifications, our method less sensitive in identifying adaptive loci in weak selection and high migration regimes. One possible resolution to this problem is to use the stationary distribution corresponding to the structured evolutionary dynamics as the process model (see Constable & McKane 2015 for an analytical solution for the stationary distribution of a structured metapopulation). Our process model may also be limited in non-equilibrium settings, such as range shifts due to rapidly changing climate or land use practices. In these cases, the quasi-stationary assumption in the process model is unlikely to hold because of transient evolutionary dynamics and eco-evolutionary feedback. Consequently, one may want to consider a probabilistic process model that jointly describes spatiotemporal changes in abundance and genetic variation (Schurr *et al*. 2012; Polechová & Barton 2015). To inform these joint process models, researchers will require temporal and spatial sampling of abundances and genomes, which can be obtained from population surveys (Pardieck *et al*. 2020) and museum collections (Payne & Sorenson 2002), respectively. These improvements in data and process models may allow biogeographers to forecast species’ response to global change using mechanistic models of ecology and evolution.

