## Supplementary Information for "Identifying genomic adaptation to local climate using a mechanistic evolutionary model"

**Model Assumptions**

The statistical model is partitioned into three levels: data, process, and parameter models. Below, we summarize the major assumptions used to construct each level.

In the data model, we assume that the population size is large, individuals are sampled randomly, and the sequenced SNPs are biallelic and independent. Next, we use the stationary distribution corresponding to an island metapopulation as our process model. This process model assumes constant (and large) effective population across demes. The genetic variation (quantified by allele frequencies) in each deme changes due to four evolutionary forces: mutation, migration, drift, and selection. Migration is assumed to be symmetric and equal for every pair of demes. We assume a linear relationship between selection and environmental variables. A biogeographer may choose alternative functional forms to represent the relationship between environmental variables and selection coefficient based on the prior scientific work. Finally, the process model assumes that at the time of sampling, allele frequencies are in quasi-equilibrium with climate. In the parameter model, we use regularized priors to induce sparsity in sensitivity coefficients. The sparsity allows us to statistically represent our prior knowledge that most of the genetic variation in the genomes stems from neutral processes.

### Non-identifiability of $\alpha_l$ and $\mu_l$

In the main text, we argued that $\mu_l$ and $\alpha_l$ are non-identifiable. Here, we demonstrate how this non-identifiability arises using both theory and simulations. First, let us consider the time-dependent changes in the deterministic part of the Fokker-Planck equation (4):

$$\frac{dp_{lk}}{dt'} = 4N_e M(p_{lk}) = 4N_e\big(s_{lk}p_{lk}q_{lk} + u_{2l}q_{lk} - u_{1l}p_{lk} + m(\overline{p_l} - p_{lk})\big),$$

where $t = 4N_e t'$. The equilibrium solution of the above differential equation (i.e., $p_{lk}^*$ for which $M(p_{lk}^*) = 0$) roughly corresponds to the mean of the stationary distribution in Eq. 11. By applying perturbation theorem, we can show the solution of $M(p_{lk}^*) = 0$ up to first order in $\tilde{s}_{lk}$ is given by

$$p_{lk}^* = \text{logit}^{-1}\big(\text{logit}(\mu_l) + \tilde{s}_{lk}\kappa_l^{-1}\big)$$

$$= \text{logit}^{-1}\left(\text{logit}(\mu_l) + \alpha_l\kappa_l^{-1} + \sum_{j=1}^{E}\kappa_l^{-1}\beta_{lj}e_{jk}\right),$$

where $\tilde{s}_{lk} = 4N_e s_{lk}$, $\mu_l = (u_{2l} + m\overline{p_l})/(u_{1l} + u_{2l} + m)$, and $\kappa_l = 4N_e(u_{1l} + u_{2l} + m)$. The first-order approximation of $p_{lk}^*$ is expressed in terms of the sum logit transformation on $\mu_l$ and a scalar multiple of $\alpha_l$, which suggests that $\mu_l$ and $\alpha_l$ are non-identifiable. However, $\beta_{lj}$ are identifiable when appropriately regularized. Next, we numerically show non-identifiability by simulating samples using the probability distribution,

$$p_{lk} \sim \frac{e^{\tilde{s}_{lk}p_{lk}}}{{}_1F_1(\mu_l\kappa_l, \kappa_l, \tilde{s}_{lk})}\text{Beta}(\mu_l, \kappa_l),$$

with parameters $\mu_l = 0.7$, $\kappa_l = 8$, $\alpha_l = 8$, and $\beta_{lj} = 0$. Then we use Stan to fit these samples to recover the parameters. We used priors $\mu_l \sim \text{Uniform}(0, 1)$, $\kappa_l \sim \text{exponential}(0.5)$, and $\alpha_l \sim \text{Normal}(0, 5)$. The pairs plot of $\mu_l$ and $\alpha_l$ (Fig. S1) shows that the posterior samples lie on an extended surface (off-diagonal plot) and the posterior distribution of $\alpha_l$ is roughly similar to its prior distribution. This suggests that $\mu_l$ and $\alpha_l$ are non-identifiable parameters.

**Parameter used to generate synthetic datasets**

To conduct model testing, we generated synthetic SNP datasets of 1000 loci per individual. The effective population size, $N_e$, in each deme (where Willow Flycatchers were sampled) is set to $100$. All loci are assigned $\alpha_l$ drawn from Normal(0, 1.25e-3). We randomly select 15 adaptive loci and pair them randomly with an environmental variable. We then assign these 15 loci $|\beta_{lj}| = 0.003$ with a randomly selected positive and negative sign. We calculate the selection coefficient in each deme using $\tilde{s}_{lk} = \alpha_l + \beta_{lj} e_j$, where $e_j$ is the standardized $j$th adaptive environmental variable. Next, we calculate standardized geodesic distances ($d_{ij}$) between every pair of demes by dividing geodesic distances by the standard deviation of all pairwise distances between demes. Next, we simulate Equation (17) for t = 200 to obtain samples from the stationary distribution. Next, we generate synthetic genotypes ($g_{ilk}$) by sampling from Binomial(2, $p_{lk}$). In each deme, we simulated genotypes for the same number of individuals as in the real data for Willow Flycatcher. We repeat this simulation 90 times to generate independent synthetic datasets representing nine evolutionary regimes—low, medium, and high mutation and migration rates (10 datasets per regime). For mutation, we use $u_{1l} = u_{2l}$ = 0.002, 0.003, and 0.004 as low, medium, and high mutation rates. For migration, we use $\rho$ = 7, 6.5, and 6 as low, medium, and high migration rates. Finally, these synthetic datasets were used as inputs to our statistical model and LFMM to calculate false negative and false positive rates (see Table S3-S6). In Fig. 4A, we used $u_{1l} = u_{2l}$ = 0.002 (low mutation) and $\rho$ = 6.5 (medium migration).

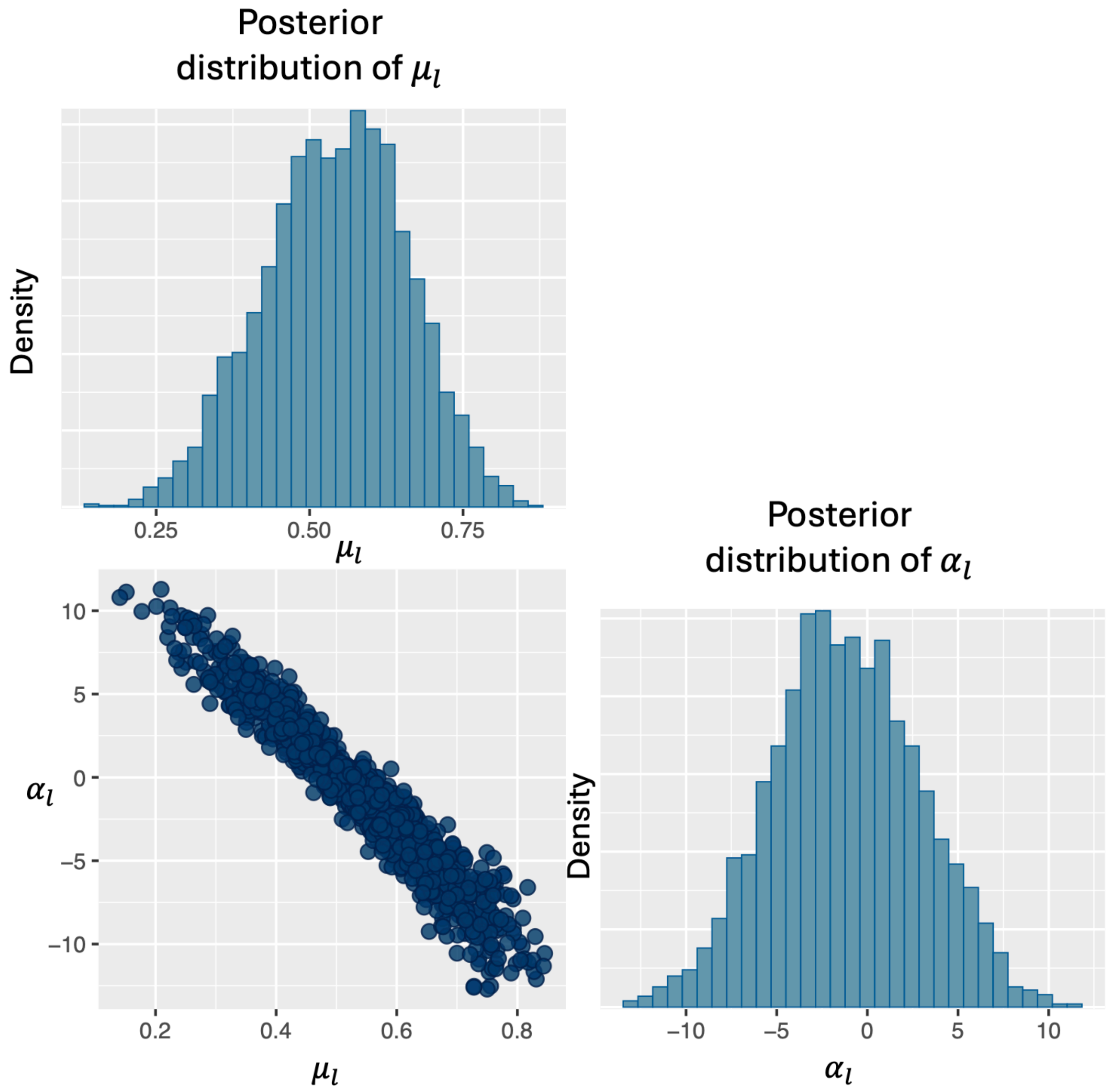


**Figure S1**: The plots show posterior samples of $\mu_l$ and $\alpha_l$. The diagonal plots are posterior distribution of $\mu_l$ and $\alpha_l$, and the off-diagonal plot shows the relationship between the posterior samples of $\mu_l$ and $\alpha_l$. The off-diagonal plot show that posterior samples lie on an extended surface, indicating that $\mu_l$ and $\alpha_l$ are non-identifiable parameters.

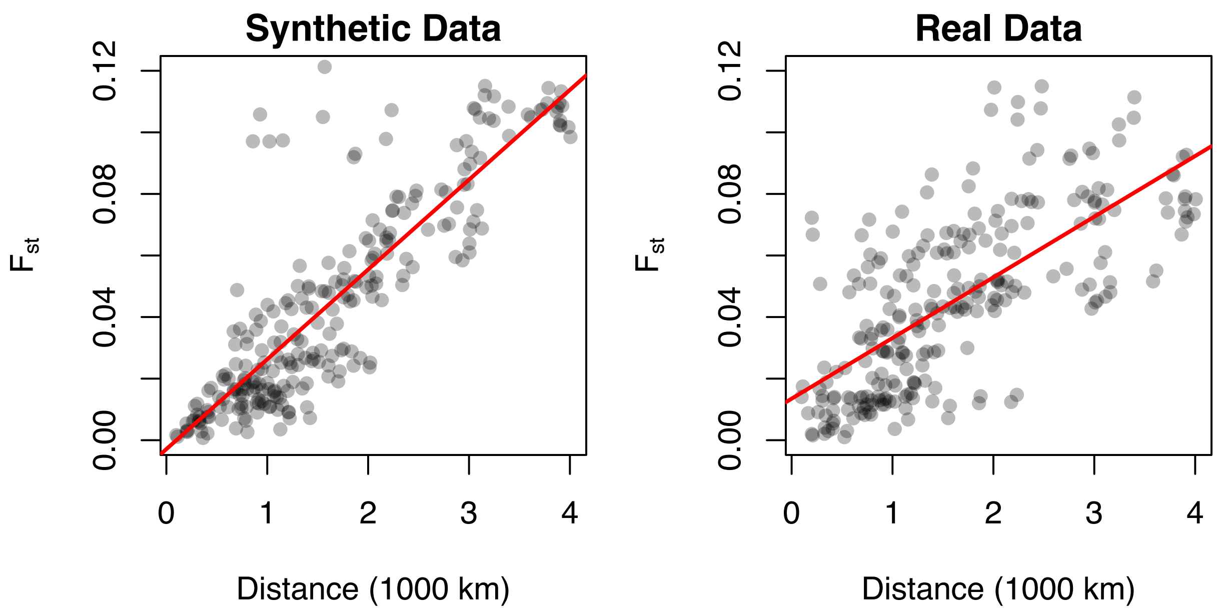


**Figure S2:** The plots show genetic differentiation (measured by $F_{st}$) between demes in synthetic and real data. In both cases, populations show signature of isolation-by-distance.

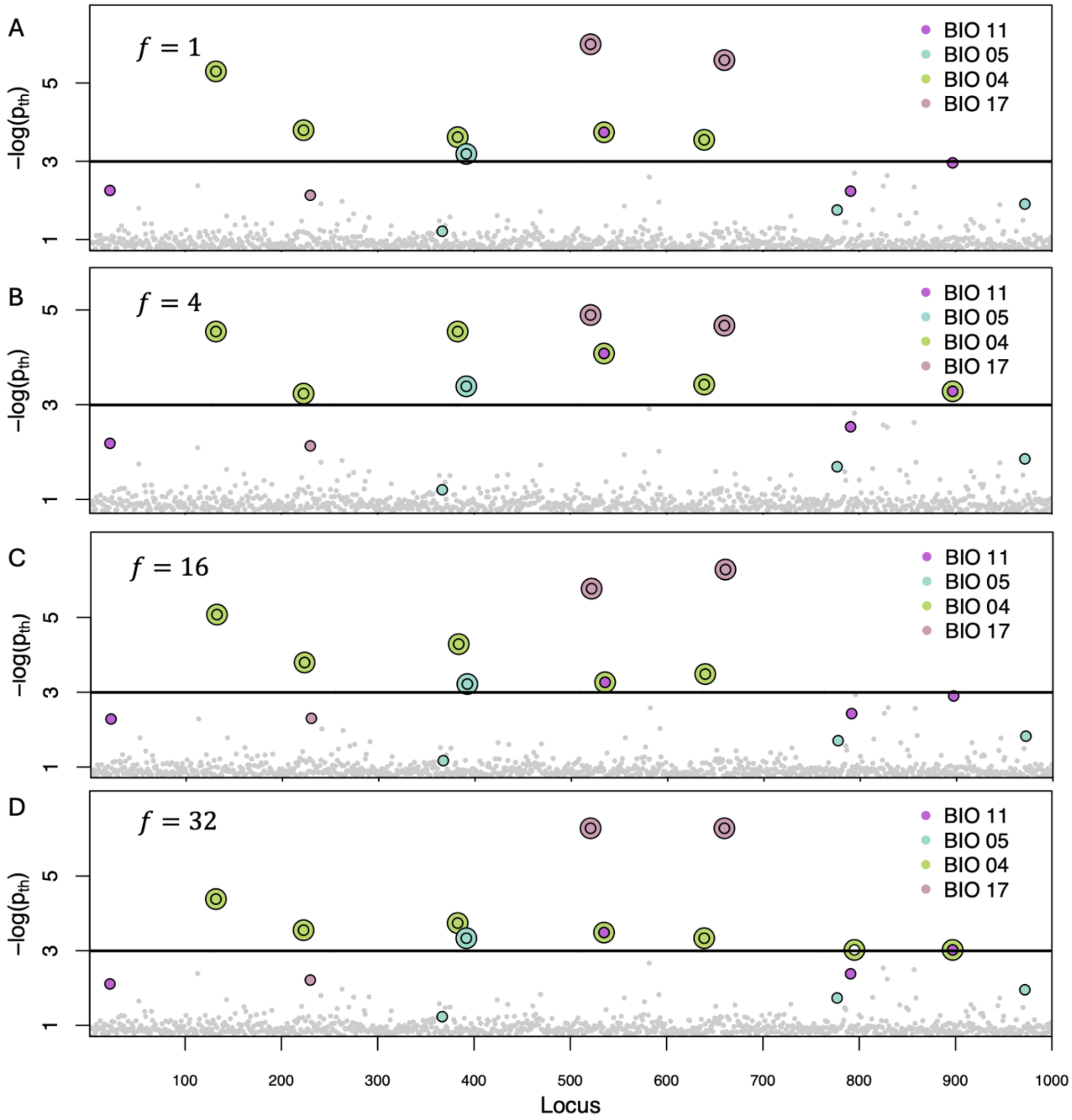


**Figure S3:** Manhattan plot showing negative log probability ($-\log(p_{th})$) that the posterior distribution of sensitivity coefficients ($\beta_{li}$) includes zero for synthetic data. This is the same dataset which was used in Fig. 4A, but here we use different values of $f$ to test the robustness of our statistical method to our choice of priors for $\tau$. We denote points above the black line using an outer ring and a center. The color of the center (ring) corresponds to the true (statistically inferred) environmental variable responsible for local adaptation.

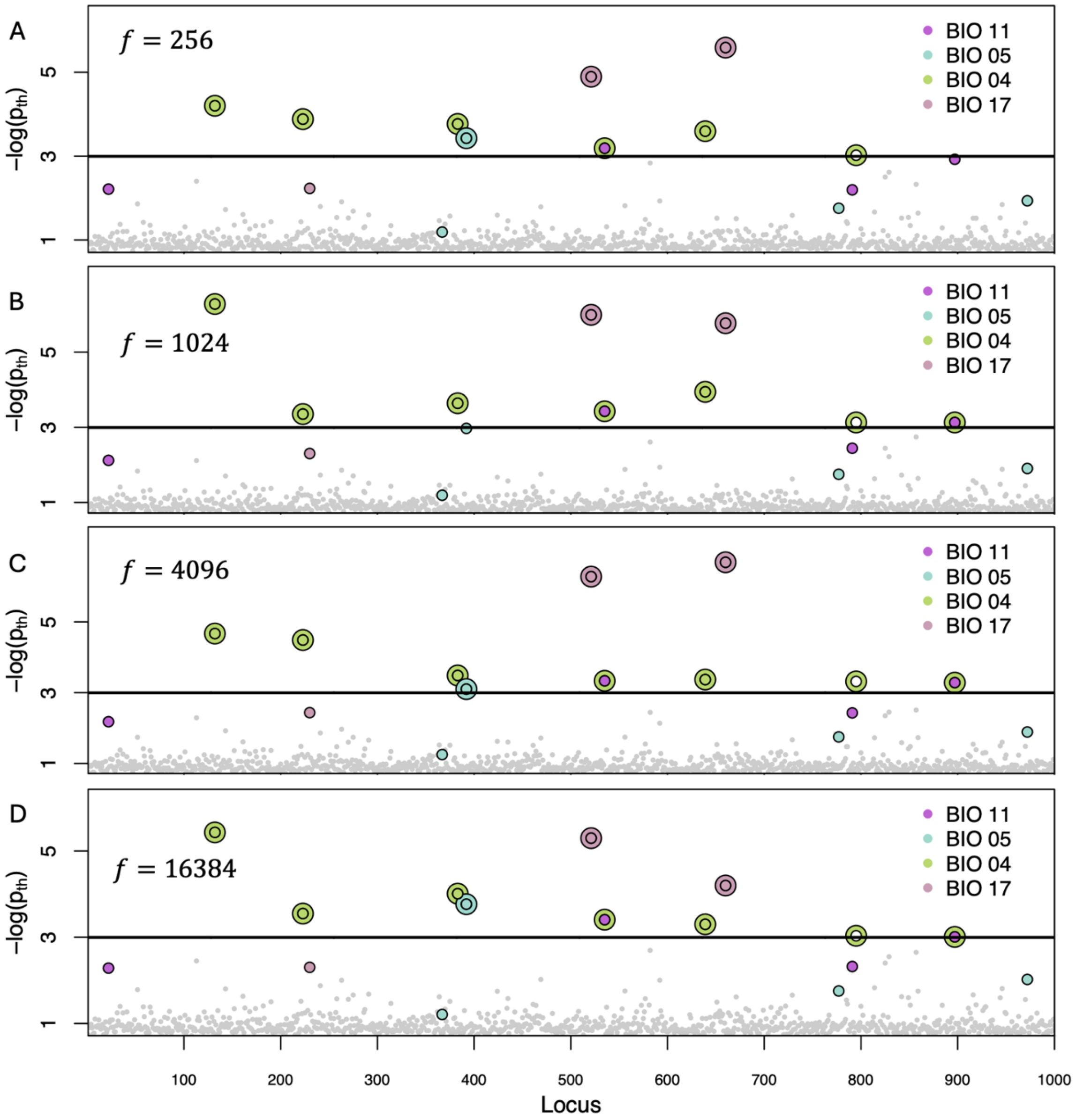


**Figure S4:** Manhattan plot showing negative log probability ($-\log(p_{th})$) that the posterior distribution of sensitivity coefficients ($\beta_{li}$) includes zero for synthetic data. This is the same dataset which was used in Fig. 4A, but here we use different values of $f$ to test the robustness of our statistical method to our choice of priors for $\tau$. We denote points above the black line using an outer ring and a center. The color of the center (ring) corresponds to the true (statistically inferred) environmental variable responsible for local adaptation.

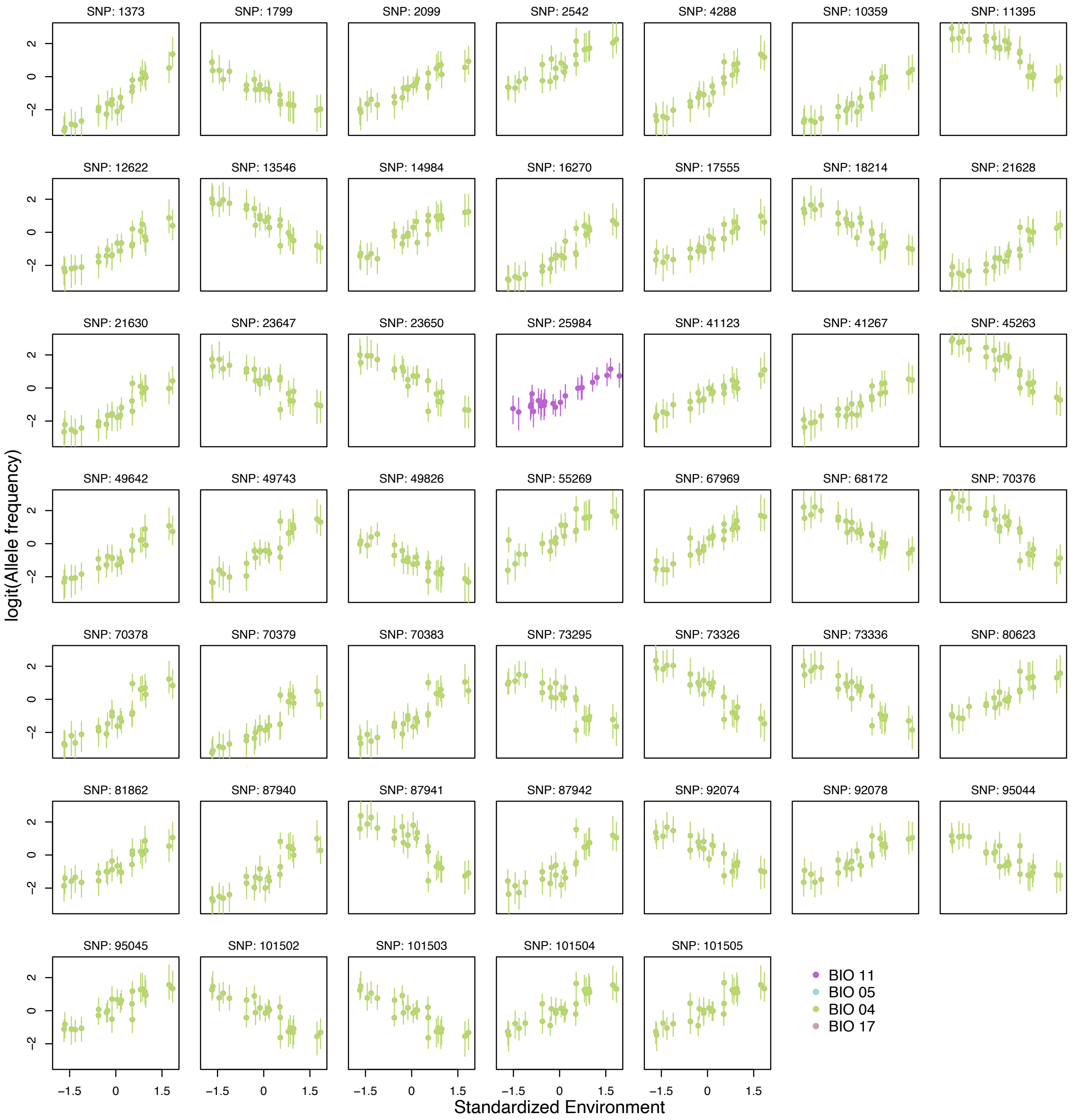


**Figure S5**: Relationship between the logit transformation of allele frequency of statistically significant loci in Figure 4B and their corresponding environmental variable.

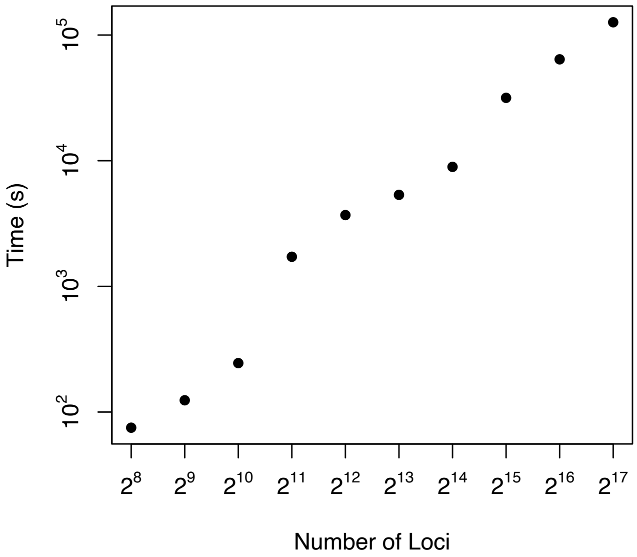


**Figure S6:** Estimated simulation time to sample posterior distribution of sensitivity coefficients as a function of the number of loci analyzed.

| Dispersal / Mutation | Low | Medium | High |
|---|---|---|---|
| Low | 34.00 | 56.67 | 80.00 |
| Medium | 55.67 | 74.00 | 87.33 |
| High | 69.33 | 86.00 | 98.00 |

**Table S1**: False negative rates in high, medium, and low migration and mutation regimes. The synthetic datasets were analyzed using the statistical model presented in this paper.

| Dispersal / Mutation | Low | Medium | High |
|---|---|---|---|
| Low | 0.41 | 0.05 | 0.01 |
| Medium | 0.03 | 0.00 | 0.00 |
| High | 0.00 | 0.00 | 0.00 |

**Table S2:** False positive rates in high, medium, and low migration and mutation regimes. The synthetic datasets were analyzed using the statistical model presented in this paper.

| Dispersal / Mutation | Low | Medium | High |
|---|---|---|---|
| Low | 99.33 | 99.33 | 99.33 |
| Medium | 98.00 | 97.33 | 98.67 |
| High | 100.00 | 99.33 | 99.33 |

**Table S3**: False negative rates in high, medium, and low migration and mutation regimes. The synthetic datasets were analyzed using LFMM with individuals' genotypes as input.

| Dispersal / Mutation | Low | Medium | High |
|---|---|---|---|
| Low | 0.00 | 0.00 | 0.00 |
| Medium | 0.01 | 0.00 | 0.00 |
| High | 0.00 | 0.01 | 0.01 |

**Table S4**: False positive rates in high, medium, and low migration and mutation regimes. The synthetic datasets were analyzed using LFMM with individuals' genotypes as input.

| Dispersal / Mutation | Low | Medium | High |
|---|---|---|---|
| Low | 100.00 | 99.33 | 99.33 |
| Medium | 98.00 | 99.33 | 98.67 |
| High | 100.00 | 98.67 | 96.66 |

**Table S5**: False negative rates in high, medium, and low migration and mutation regimes. The synthetic datasets were analyzed using LFMM with raw allele frequencies as input.

| Dispersal / Mutation | Low | Medium | High |
|---|---|---|---|
| Low | 0.13 | 0.17 | 0.13 |
| Medium | 0.13 | 0.17 | 0.12 |
| High | 0.21 | 0.18 | 0.14 |

**Table S6**: False positive rates in high, medium, and low migration and mutation regimes. The synthetic datasets were analyzed using LFMM with raw allele frequencies as input.

| Gene | Broad Function | Gene | Broad Function |
|---|---|---|---|
| EDIL3 | Egg shell mineralization | PITPNM3 | Vision |
| FCHSD1 | Enable lipid binding activity | HBS1L | Control hemoglobin level |
| CELF6 | Slicing and editing mRNA | CHST11 | Immune response |
| ATG5 | Immune system | PARP9 | Immune system |
| SOX14 | Cell proliferation and neuronal differentiation | ATP11C | Development of B and red blood cells |
| AKAP6 | Skeletal myoblast differentiation and muscle regeneration | GAS7 | Maturation of cerebellar neurons |
| SMTNL2 | Enable protein phosphatase 1 and tropomyosin binding activity. | KCNF1 | Modulate vocal motor function |
| UHRF2 | Cell cycle regulation | GSG1L2 | Plasma membrane |
| PCDH1 | Feather bud | RAD23B | Nucleotide excision repair |
| LOC114065648 | Uncharacterized | SPINT1 | Uncharacterized |
| GRIK2 | Uncharacterized | LOC114056459 | Uncharacterized |
| SALL1 | Uncharacterized | CCDC61 | Uncharacterized |
| LOC114061458 | Uncharacterized | LOC114067681 | Uncharacterized |
| LOC114062095 | Uncharacterized | TMCO4 | Uncharacterized |
| LOC114062096 | Uncharacterized | LOC114071423 | Uncharacterized |

**Table S7:** Broad function of 30 genes that are present within the 25kb distance from loci that have statistically significant sensitivity coefficient.
